# An ultra-low frequency spike timing dependent plasticity-based approach for treating alcohol use disorder

**DOI:** 10.1101/2021.09.16.460673

**Authors:** Anders J. Asp, Suelen Lucio Boschen, J. Luis Lujan

## Abstract

Alcohol use disorder (AUD) is a chronic relapsing brain disorder characterized by an impaired ability to stop or control alcohol consumption despite adverse social, occupational, or health consequences. AUD affects nearly one-third of adults at some point during their lives, with an associated cost of approximately $249 billion annually in the U.S. alone. The effects of alcohol consumption are expected to increase significantly during the COVID-19 pandemic, with alcohol sales increased by approximately 54%, potentially exacerbating health concerns and risk-taking behaviors. Unfortunately, existing pharmacological and behavioral therapies for AUD have historically been associated with poor success rates, with approximately 40% of individuals relapsing within three years of treatment.

Pre-clinical studies have shown that chronic alcohol consumption leads to significant changes in synaptic function within the dorsal medial striatum (DMS), one of the brain regions associated with AUD and responsible for mediating goal-directed behavior. Specifically, chronic alcohol consumption has been associated with hyperactivity of dopamine receptor 1 (D1) medium spiny neurons (MSN) and hypoactivity of dopamine receptor 2 (D1) MSNs within the DMS. Optogenetic, chemogenetic, and transgenic approaches have demonstrated that reducing the D1/D2 MSN signaling imbalance decreases alcohol self-administration in rodent models of AUD. However, these approaches cannot be studied clinically at this time.

Here, we present an electrical stimulation alternative that uses ultra-low (<=1Hz) frequency (ULF) spike-timing dependent plasticity (STDP) to reduce DMS D1/D2 MSN signaling imbalances by stimulating D1-MSN afferents into the GPi and ACC glutamatergic projections to the DMS in a time-locked stimulation sequence. Our data suggest that GPi/ACC ULF-STDP selectively decreases DMS D1-MSN hyperactivity leading to reduced alcohol consumption without evoking undesired affective behaviors in a two-bottle choice mouse model of AUD.

## Introduction

Alcohol Use Disorder (AUD) is defined as a chronic relapsing brain disorder characterized by an impaired ability to stop or control alcohol consumption despite adverse social, occupational, or health consequences ^31^ and is one of the most common psychiatric disorders. AUD affects nearly one-third of U.S. adults at some point during their lives and carries a financial burden greater than $249 billion annually ^1^. Furthermore, it leads to approximately 88,000 preventable deaths in the U.S. each year ^32^. Existent pharmacological and behavioral therapies for AUD have had poor success rates ^2,3^, with approximately 40% of individuals relapsing within three years of treatment ^33^.

It has been shown that chronic alcohol consumption leads to significant changes in glutamatergic synaptic function within the dorsal medial striatum (DMS) ^13,18,20,27^, one of the brain regions associated with underlying maladaptive changes observed in AUD and responsible for mediating goal-directed behavior ^12^. The DMS receives cortical glutamatergic input from limbic regions, such as the medial prefrontal cortex and anterior cingulate cortex (ACC) ^34,35^. The DMS contains two neuronal types: 1) Dopamine receptor 1-medium spiny neurons (D1-MSNs) and 2) Dopamine receptor 2-medium spiny neurons (D2-MSNs). D1-MSNs project to the internal Globus Pallidus (GPi) through the direct pathway and their activation results in a ‘*go*’ signal to initiate behavior ^36–40^. In contrast, D2-MSNs project to the external Globus Pallidus (GPe) through the indirect pathway and their activation serves as a ‘*stop*’ signal to inhibit behavior ^36–40^. Chronic alcohol consumption has been shown to cause permanent hyperactivity of D1-MSNs and permanent hypoactivity of D2-MSNs in the DMS via maladaptive cortical glutamatergic signaling from areas such as the ACC ^13–17^. Studies have shown that reducing the ratio of DMS D1-MSN/D2-MSN signaling imbalance reduces pathological alcohol seeking behavior ^17,20–29^. Furthermore, a recent report demonstrated that inducing long-term depression (LTD) by applying a single 10-minute epoch of low frequency (1Hz) optogenetic stimulation of ACC projections to the DMS, combined with systemic D1-dopamine receptor antagonism, leads to a reduction in alcohol consumption lasting for nine days ^26^.

Data suggest that deep brain stimulation (DBS), an increasingly prevalent therapy for motor and psychiatric disorders ^41–44 6,7,11,45–48 49^, may offer therapeutic effects for the treatment of AUD ^4,6–10^. However, unlike DBS for movement disorders, there is an absence of markers of DBS efficacy for the treatment of addiction and other psychiatric conditions ^50,51^, leading to challenging selection of surgical targets and stimulation parameters. These factors, combined with nonspecific network effects of DBS ^52^, can lead to undesirable side effects and highly variable outcomes in the treatment of addiction-related behaviors such as AUD ^5,11,53,54^. Thus, re-imagining the way that DBS is administered may lead to the development of new therapeutic interventions for a wide range of neuropsychiatric and neurologic disorders. Here, we leverage neuroplasticity induction protocols capable of selectively reversing the pathophysiology underlying neuropsychiatric conditions such as AUD. ^30,55–57^. Changes in synaptic strength are evoked by leveraging spike-timing dependent plasticity (STDP) ^30,55–57^, a natural phenomenon underlying learning and memory. In STDP, bidirectional control of network gain can be achieved by repeated phase-locked activation of pre- and post-synaptic neural elements. The direction and magnitude of the spike-timing dependent synaptic modulation is governed by the relative timing between pre- and post-synaptic depolarization **(Figure 1A, 1B)**. For example, long-term potentiation (LTP) is caused when postsynaptic action potentials occur repeatedly after pre-synaptic action potentials (negative timing, STDP(-)). On the other hand, long-term depression (LTD) is caused when post-synaptic action potentials occur repeatedly prior to pre-synaptic action potentials (positive timing, STDP(+)) ^55,57^ **(Figure 1C)**. Here, we describe an electrical stimulation approach that uses ultra-low (≤1Hz) frequency spike-timing dependent plasticity (ULF-STDP)^30^ to reduce DMS D1/D2 MSN signaling imbalances by repeatedly stimulating the D1-MSN afferents into the GPi before stimulating ACC glutamatergic projections to the DMS **(Figure 1D)**. Action potentials originating in D1-MSN axons within the GPi resulting from electrical stimulation will backpropagate to their somas in the DMS^58^ and depolarize postsynaptic dendrites^59–61^. By leveraging the anatomical separation of D1-MSNs projections to the GPi via the direct pathway^36,62^, D1-MSN axons in the GPi can be depolarized via electrical stimulation without depolarizing D2-MSN axons in the GPe. In this manner, pairing phase-locked presynaptic ACC cortical stimulation with postsynaptic GPi stimulation could provide selective control of spike-timing dependent synaptic strength in D1-MSNs while avoiding changes in synaptic strength of D2-MSNs projecting to the GPe **(Figure 2)**.

**Figure 1:**
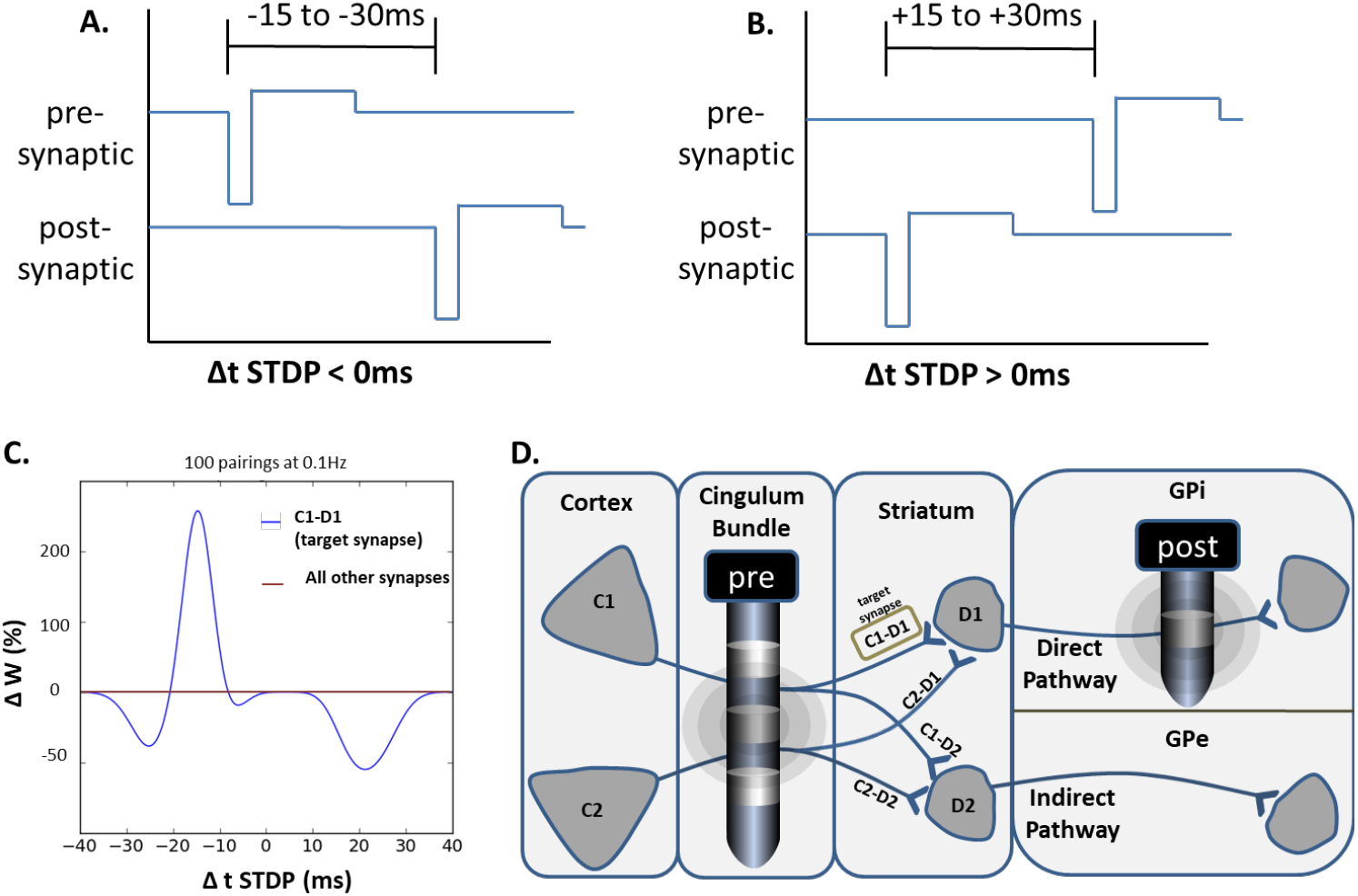
ULF-STDP(+/-) stimulation approach for bi-directional network gain control. ULF-STDP(-) **(A)** and ULF-STDP(+) **(B)** protocol schematics. Biophysical model of STDP(+) adapted from (Cui, et al., 2016)^56^ **(C)**. Schematic representation of ULF-STDP(+) protocol where an electrical stimulus is applied to cortical axons C1/C2 within a temporal window with respect to stimulation of postsynaptic axons D1/D2 **(D).** In this example, orthodromic action potential propagation from ACC axons and antidromic action potential propagation from D1 axons in GPi will converge to alter the gain of the spatially unique synapse (C1-D1) while gain of synapses out of this network are unchanged. This protocol is repeated at ≤1Hz for 10 minutes.

**Figure 2:**
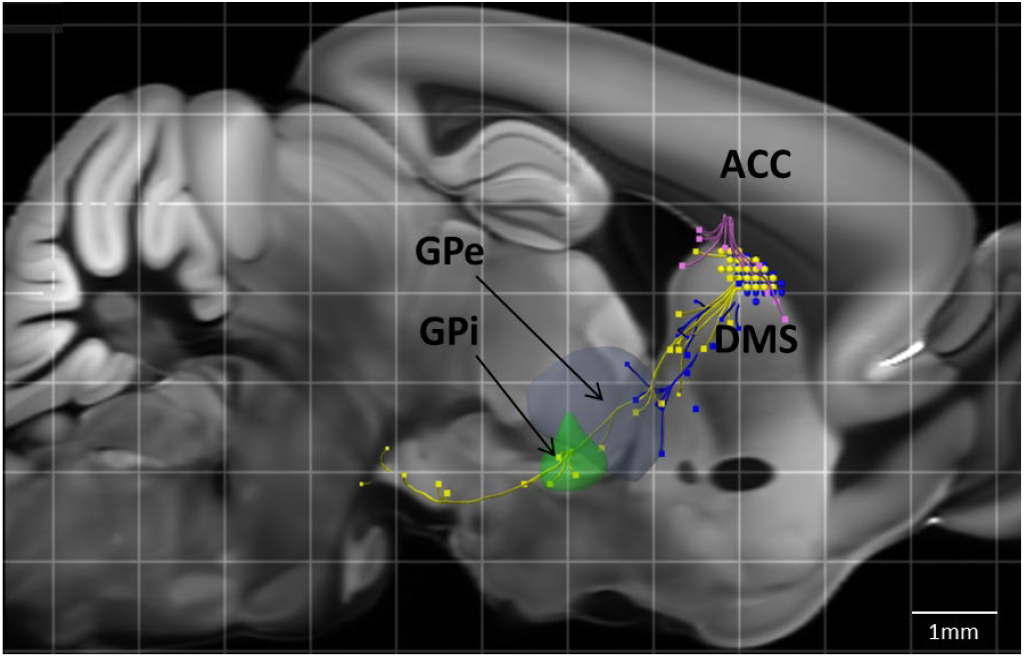
D1-MSN efferents are distinct and separable from D2-MSN efferents. D1-MSN efferents (yellow), can be targeted by placing a stimulating electrode in the GPi while avoiding D2-MSN efferents (blue). ACC efferents project to DMS (magenta). Data source: Allen Mouse Brain Connectivity Atlas (2011). Experiments 159223001 and 127224133.

## Methods

### Animals

All behavior experiments were performed with wild type C57/Bl 6J mice (Jackson Laboratory, Bar Harbor, Maine, JAX Stock No: 000664). Fluorescent reporter mouse lines Drd1a-tdTomato mice (JAX Stock No: 016204) were used for labeling D1-MSNs in patch clamp electrophysiology experiments. All laboratory procedures were reviewed and approved by the Mayo Clinic Institutional Animal Care and Use Committee (IACUC) and conform to guidelines published by the NIH Guide for the Care and Use of Laboratory Animals (Department of Health and Human Services, NIH publication No. 86-23, revised 1985). Mice were housed in plastic cages under standard 12-hour light/dark cycle and conditions (21°C, humidity 45%) with ad libitum access to food and water. Animals were acclimated for at least one weeks before use. All efforts were made to minimize both the number of mice used and any discomfort that may be experienced.

### Stereotactic surgery

All animals were induced at 4% isoflurane and maintained at 1-2% through a nose cone for the duration of the surgery. The depth of anesthesia was monitored using toe pinch and eye blink reflexes. A heating pad was used to maintain the subject’s body temperature to 37.0±0.5 °C throughout the duration of anesthesia. Analgesia was provided with buprenorphine HCL (0.05 mg/kg). Additionally, Ibuprofen was delivered in the animals’ drinking water two days prior to the surgery and no less than five days after for analgesia. Animals were monitored twice daily for five days following surgery for signs of distress and infection at the surgical site. If signs of infection or distress were observed, topical polysporin antibacterial ointment was applied to the surgical site or a veterinarian was consulted for the appropriate methods of treatment.

A stereotaxic frame (Model 1900, David Kopf Instruments, Tujunga, CA) was used for implantation of custom built bipolar 50.8 micron diameter Pt/Ir wire insulated with PTFE (A-M systems 77600, Sequim, WA) and recording Zif-Clip microwire arrays (Tucker-Davis Technologies, Alachua, FL). All surgical instruments, electrodes, and skull screws were sterilized in an autoclave or MetriCide28 (Metrex, Oragnge, CA, USA) cold sterilant. Anaesthesia was induced in mice with isoflurane 4% followed by 1-2% isoflurane for maintenance. The surgical sites on mice heads were shaved and cleaned with betadine. Then, mice were placed in the stereotaxic apparatus and secured using ear bars. An incision of approximately 1.5 cm was made in the skin over the skull with a scalpel. Hydrogen peroxide was used to clean the skull surface. Sterile screws were used to provide a strong fixation of the head cap and serve as an electrical reference and ground. Screws were secured into 0.7mm holes drilled in the skull with a trephine drill bit. An additional craniotomy, approximately 2 mm x 2 mm in size, was created over the DMS to implant the Zif-Clip microwire array in a subset of animals. The dura was removed to expose brain tissue, which was irrigated with cool saline throughout the surgery. For all surgical procedures, additional burr holes were drilled at the desired coordinates described next to create a cranial window for unilateral implantation of custom-built stimulating electrodes per the Mouse Brain in Stereotaxic Coordinates ^12^. Stimulating electrodes were placed in the GPi (AP: −1.4 mm, ML: 1.6 mm, DV: −4.5 mm from Bregma) and ACC (AP: 0.50 mm, ML: 0.7 mm, DV: −1.85 mm from Bregma). Upon insertion and fixation of the stimulating electrodes to the skull using dental cement (Henry Schein, Melville, NY), a Zif-Clip microwire array was chronically implanted into the DMS (array center: AP: 0.70 mm, ML: 1.5 mm, DV: −2.75 mm from Bregma). Kwik-Sil silicone elastomer (World Precision Instruments, Sarasota, FL, USA) was put around the Zif-Clip microwire array and stimulating electrodes to maintain a seal between the brain and dental cement described in the next step. Finally, all components were secured with Metabond dental cement (Parkell, Edgewood, NY, USA). After the electrode implantation surgery, the animals were housed individually for one week to allow sufficient time for recovery.

### Electrical stimulation parameters

Electrical stimulation was delivered through custom-built bipolar parallel chronically indwelling stimulating electrodes via a IZ2M-64 microstimulator programmed in the Synapse software (Tucker-Davis Technologies, Alachua, FL). The ultra-low frequency spike-timing dependent plasticity stimulation (ULF-STDP) protocol consists of a charge-balanced, biphasic, cathodic-leading, 250 μA current stimulation with a pulse duration of 90μs delivered to the GPi and ACC with 18ms between GPi and ACC stimulation. GPi-ACC stimulation pairing was delivered at 1Hz for 10 minutes. ULF-STDP (+) is defined as postsynaptic GPi stimulation 18 ms prior to presynaptic ACC stimulation and ULF-STDP (-) is defined as presynaptic ACC stimulation 18 ms prior to postsynaptic GP stimulation. Current was delivered at a density of 941 μC/cm^2 such that Shannon safety criteria are satisfied to avoid tissue damage^13^.

### Two-bottle choice alcohol self-administration AUD model

A standard two-bottle choice paradigm was used to assess alcohol consumption^63^. Briefly, two bottles were presented daily: one containing water and the other alcohol ^64^. The position of the alcohol and water bottles was swapped daily to reduce confounds produced by location preference. For the first week, alcohol concentrations were increased every other day from 3% to 6% to 10% v/v. For the remaining duration of the experiment, mice were presented with one bottle of water and another bottle with 10% alcohol. For three alcohol exposure days prior to ULF-STDP delivery, the animals were tethered to the stimulating system to allow habituation to the procedure. At day 25, animals were stimulated with the ULF-STDP protocol described in the methods section under *Electrical stimulation parameters* if mean and standard error of alcohol consumption (calculated in g/kg of body mass at be beginning of the night cycle) was less than 20% over a 3-day period after at least two weeks of alcohol exposure. The mass of each mouse, the mass of alcohol consumed, and the mass of water consumed was assessed daily. Additionally, an IR beam break sensor was placed in front of the sippers to measure the total time spent interacting with the sipper.

### In vivo electrophysiology

Previously described standard methods were followed for *in vivo* electrophysiological recordings of MSNs within the DMS ^59^. Briefly, mice were surgically implanted with chronic indwelling Pt/Ir stimulating electrodes in the GPi and ACC along with a custom 32-channel ZIF-Clip microwire array described in the *Stereotactic Surgery* section. Mice were given one week to recover from surgery prior to recordings during unrestrained locomotion. Electrophysiological recordings were performed in three stages: alcohol naïve, following chronic alcohol exposure, and following ULF-STDP stimulation. Multichannel extracellular recordings were performed at 50 kHz and band-pass filtered from 300 Hz to 5 kHz prior to spike thresholding and sorted using principal component analysis (PCA) via the Synapse software (TDT). Mean event rates over 10-minute periods were compared before and after ULF-STDP application using a Student’s unpaired t-test (p<0.05).

### *Ex vivo* whole-cell patch clamp electrophysiology

We followed standard procedures for slice preparation and *ex vivo* whole cell electrophysiology to measure D1-MSN activity ^27,65,66^. Five week-old alcohol-naiive Drd1a-tdTomato mice (JAX Stock No: 016204) received ULF-STDP(+) *in vivo* under 1-2% isoflurane anesthesisia 30 minutes prior to euthanasia for whole cell patch clamp electrophysiology experiments. Briefly, cells were clamped at −75 mV in the presence of lidocaine (0.7 mM) to record spontaneous mEPSCs. The mEPSC amplitude and frequency were compared between animals which received ULF-STDP(+) and stimulation naiive animals. Statistical significance (p<0.05) was determined via unpaired Student’s t-tests. D1-MSNs were identified by fluorescence microscopy in Drd1a-tdTomato mice.

### Euthanasia and Histology

At the end of the experiments, animals were euthanized by a lethal overdose of Pentobarbital (i.p.), in accordance with the Panel on Euthanasia of the American Veterinary Medical Association (A.V.M.A.). Mice were transcardially perfused with 4% paraformaldehyde in phosphate buffered saline (PBS). Mice brains were removed and dehydrated in 30% sucrose prior to flash-freezing with dry ice and slicing into 40 μm-thick coronal sections using a sliding microtome (Leica Biosystems, Buffalo Grove, IL). Slices were mounted and counterstained with DAPI (Vector Laboratories, Burlingame, CA, USA) and visualized using bright field and fluorescence microscopy (Carl Zeiss LSM780 or Nikon Eclipse FN1) to verify electrode location. Animals were included in behavioral and electrophysiological analysis only if at least one of each bipolar electrode tips were located in both the ACC and GPi as defined by atlas boundaries.^12^

### Rigor and reproducibility

The observed effect size of ULF-STDP in reducing alcohol consumption was *d_Cohen_* = 0.603, in which 75% of the mice (15 out of 20 mice) developed high alcohol preference in the two-bottle choice model of AUD. Animal numbers for each cohort in each experiment were determined based on an 80% statistical power, 95% confidence interval, and effect size of 0.603. Data collected were tested for normality with the Kolmogorov-Smirnov test. Comparisons of non-normal distributions were performed using nonparametric tests, substituting the Mann-Whitney U-test in place of the unpaired Student’s t-test.

## Results

### ULF-STDP allows bidirectional control of alcohol consumption in a two-bottle choice model of AUD

Our initial goal was to determine if ULF-STDP(+) decreases alcohol consumption in a two-bottle choice mouse model of AUD. Mice implanted with stimulating electrodes in the ACC and GPi reached stable consumption of 10% v/v alcohol and were separated into two groups, a high alcohol preference group (>60% preference, n=15) and low alcohol preference group (<60% preference, n=5) **(Figure 3A)**. ULF-STDP(+) reduced alcohol consumption in the high alcohol preference group with a mean of differences reduction of 3.2 g/kg/24hr (n=15, p<0.05) **(Figures 3A, B)**. Conversely, ULF-STDP(-) increased alcohol consumption in the low alcohol preference group with a mean of differences increase of 5.2 g/kg/24hr (n=5, p<0.05) **(Figures 3A, C)**. ULF-STDP(+) did not affect food **(Figure 3D)** or water **(Figure 3E)** consumption in the high alcohol preference group (n=5, p<0.05). ULF-STDP(+) reduced time drinking alcohol for 170 minutes **(Figure 3F)**, but this effect was absent during the period of 360-490 minutes after stimulation **(Figure 3G)**, suggesting reductions in alcohol consumption may be transient in nature. Importantly, alcohol consumption in the two-bottle choice model was not altered when the ULF-STDP(+) protocol was applied to only the ACC or only the GPi electrode (**Figure S1).**

**Figure 3:**
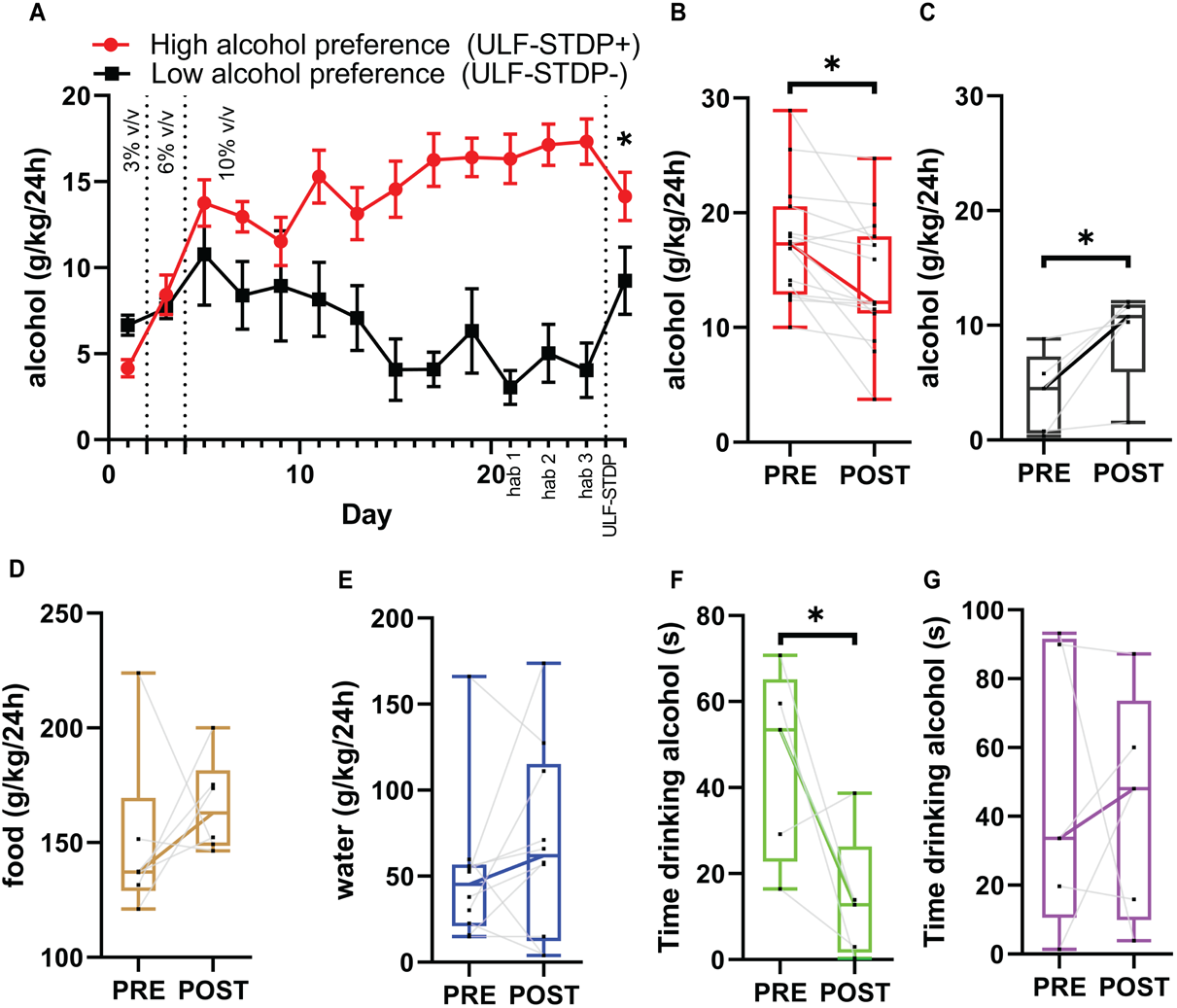
ULF-STDP allows bidirectional control of alcohol consumption in a two-bottle choice model of AUD without influencing other affective behaviors such as food and water consumption. ULF-STDP(+) reduces alcohol consumption in mice displaying high alcohol preference, while ULF-STDP(-) increases alcohol consumption in mice displaying low alcohol preference **(A)**. Alcohol consumption in g/kg/24h period before ULF-STDP(+/-) (PRE) and 24 hours after ULF-STDP(+/-) (POST) for both the ULF-STDP(+) **(B)** and ULF-STDP(-) **(C)** protocols. Food **(D)** and water **(E)** consumption (g/kg/24h). Time drinking alcohol as determined by an IR beam break sensor placed in front of the alcohol sipper for the first 170 minutes of the night cycle on the day before (PRE) and after (POST) ULF-STDP(+) **(F)** and 360-490 min of the night cycle on the day before (PRE) and after (POST) ULF-STDP(+) **(G)**. *PRE* indicates the 24 hours before ULF-STDP(+/-) and *POST* indicates the 24 hours after ULF-STDP(+/-). Statistical significance of comparison of alcohol consumption immediately before and after ULF-STDP was determined using Student’s t-test, p<0.05 (n=15). Error bars represent standard error. Box and whisker plots in B-G represent median, 95% CI, and max/minimum where black dots and gray lines represent individual animals.

### ULF-STDP(+) reduces DMS mean firing rate

Our next goal was to characterize if ULF-STDP(+) alters MSN firing patterns within the DMS underlying behavioral changes. Single units were recorded i n the DMS from a chronically indwelling microwire arrays (Figure 4A, B) in high alcohol preference mice before and after ULF-STDP (+). The median MSN firing rate in high alcohol preferring rate calculated over a 10-minute period was reduced from 0.99Hz to 0.76Hz after ULF-STDP(+) **(Figure 4C)** (n=3 mice, n=65 cells, p<0.05). Only MSN units were used in firing rate calculations. MSNs **(Figure 4D, left)** were distinct and separable from non-MSN units **(Figure 4D, right)** in the DMS, as determined by PCA. The reduction in MSN firing rate is shown in the representative DMS microwire array channel of a high alcohol preference animal before **(Figure 4E, top)** and after **(Figure 4E, bottom)** ULF-STDP(+). Microwire array impedances were stable over the course of the experiment (Figure S1). Additionally, there were no differences in spectral band power before and after ULF-STDP (Figure S2).

**Figure 4:**
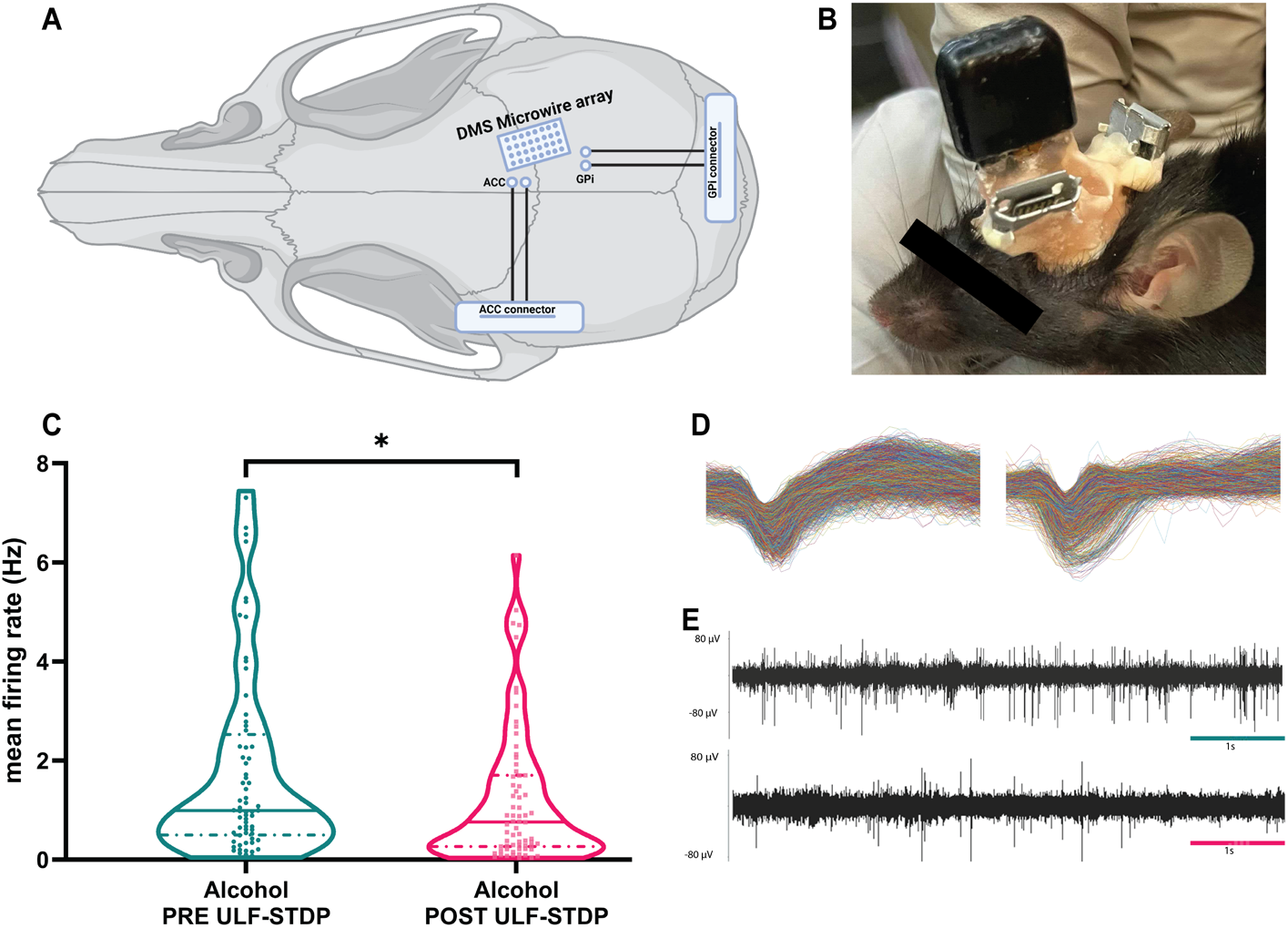
Electrophysiological DMS activity in AUD mice before and after ULF-STDP(+). Schematic **(A)** and actual **(B)** placement of recording microwire electrode array (MEA) and stimulating electrodes in the DMS, ACC, and GPi, respectively. Mean firing rate of DMS MSNs from alcohol-exposed animals before and after ULF-STDP(+) **(C)**. Representative MSN (left) and non-MSN (right) **(D)**. Representative filtered traces from microwire array recordings in DMS of alcohol-exposed animals before (top) and after (bottom) ULF-STDP(+) **(I)**. Statistical significance determined using two-tailed Mann-Whitney test, *p*<0.05.

### ULF-STDP(+) reduces evoked multiunit potentials in DMS in vivo

The mechanism of action of ULF-STDP is predicated on altering the strength of synaptic connections between the cortex and striatum. Next, we tested if ULF-STDP(+) specifically reduced evoked corticostriatal multiunit potentials in the DMS of anesthetized mice. Representative traces of multiunit potentials evoked from GPi stimulation and ACC stimulation are shown in Figures 5A and 5B. ULF-STDP(+) reduced the peak evoked multiunit potential from ACC stimulation with a mean of differences of 17.97μV before and after ULF-STDP(+), but no change was observed in responses evoked from GPi stimulation **(Figure 5C)** (n=3 mice, n=21 channels, p<0.05). Latency from stimulation to peak evoked multiunit response did not differ between stimulation location and was not affected by ULF-STDP(+) **(Figure 5D)**.

**Figure 5:**
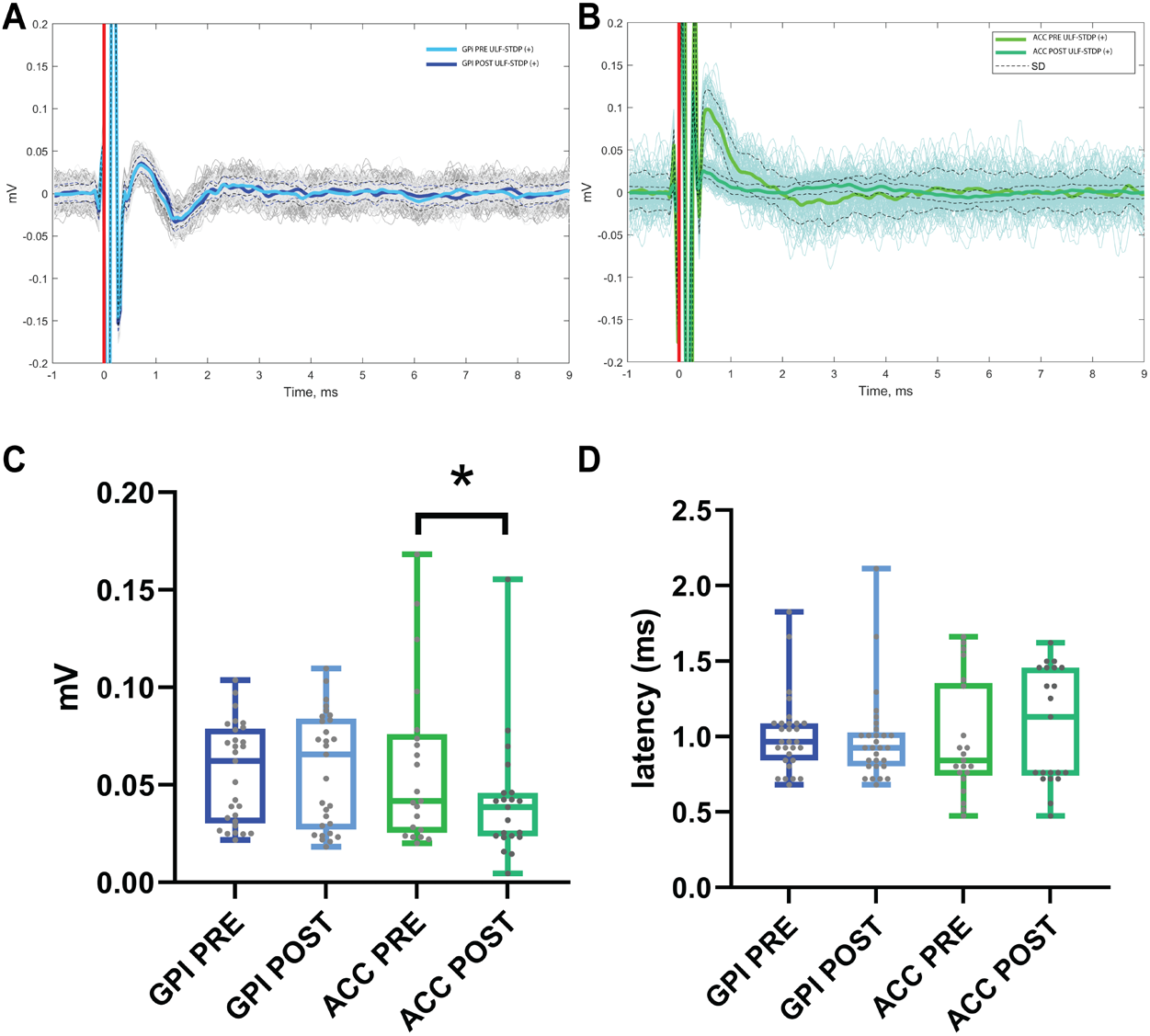
ULF-STDP(+) reduces evoked multi-unit potentials in DMS *in vivo*. Example of acute microwire array recordings of multiunit responses evoked by stimulation of the GPi **(A)** or ACC **(B)** stimulation before and after ULF-STDP(+). Peak evoked amplitudes **(C)** and latencies to peak of evoked potentials are described **(D)**. Statistical significance determined by paired Student’s t-test (n=3 mice, n=21 channels, p<0.05).

### ULF-STDP(+) reduces synaptic strength of D1-MSNs of the DMS ex vivo

The D1-MSNs in the DMS shows increased glutamatergic transmission after chronic exposure to alcohol ^13–17^. Reducing the synaptic strength of DMS D1-MSNs also reduces alcohol consumption ^20^. Therefore, we tested if ULF-STDP(+) reduces synaptic strength of DMS D1-MSNs by comparing *ex vivo* D1-MSN miniature excitatory postsynaptic current (mEPSC) amplitude and frequency in the DMS of stimulation naïve animals to those which receive ULF-STDP(+) *in vivo*.Example traces of D1-MSNs in the DMS identified with a positive Td-Tomato reporter are shown in **Figure 6A**. ULF-STDP(+) decreased D1-MSN mEPSC amplitude **(Figures 6B, C)**, but not the frequency **(Figure 6D, E)** (ULF-STDP(+) n=1 mouse, n=3 cells; naiive n=5 mice, n=1-3 cells/mouse, p<0.05).

**Figure 6:**
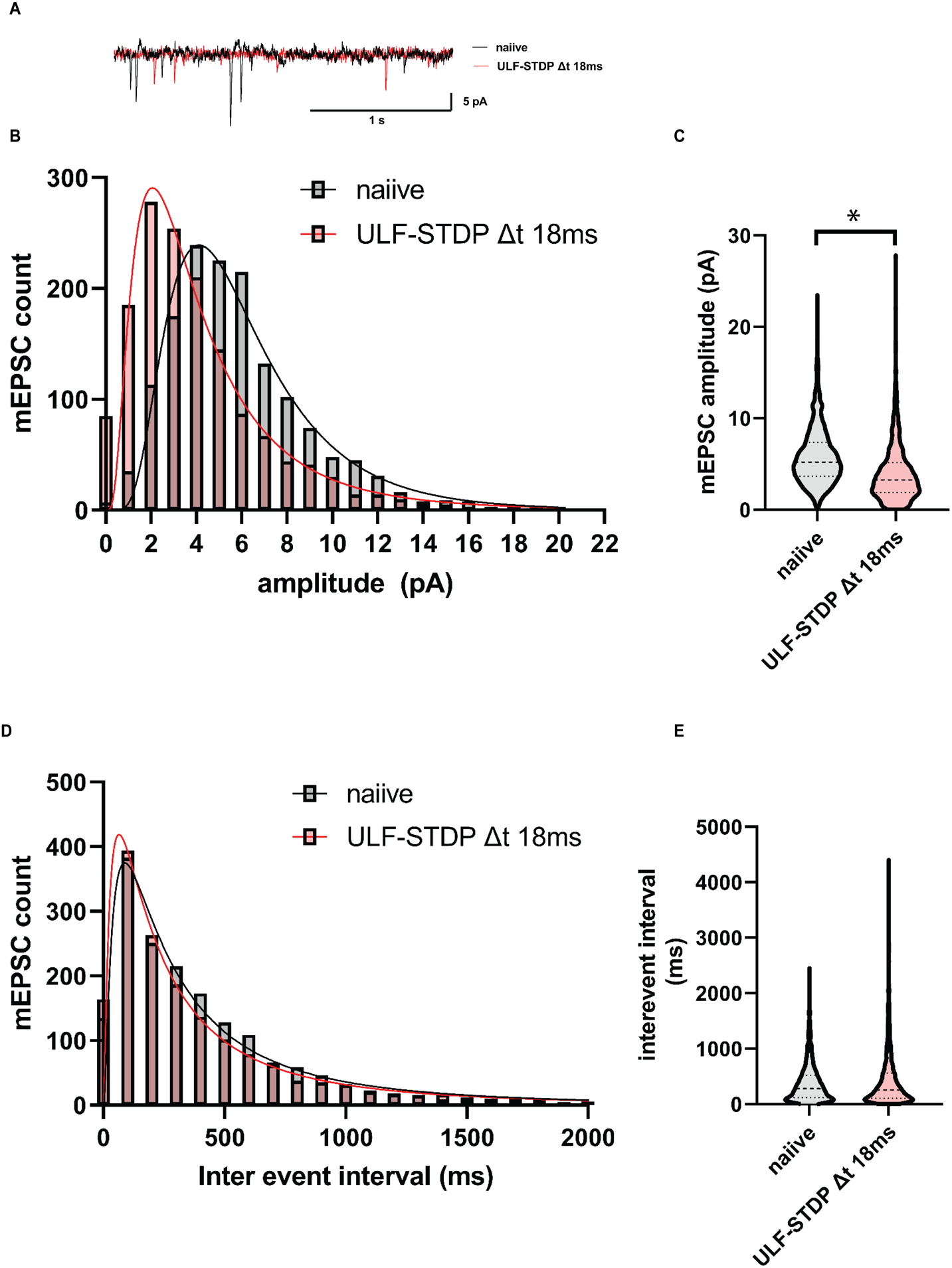
GPi/ACC ULF-STDP(+) decreases mEPSC amplitude, but not latency. Example current traces from DMS D1-MSNs clamped at −75mV of naiive mice (black) and mice treated with ULF-STDP(+) *in vivo* **(A)** Histogram showing mEPSC amplitude distribution **(B)** and violin plot of mEPSC amplitude **(C).** Histogram showing mEPSC inter-event interval distribution **(D)** and violin plot of mEPSC inter-event interval **(E).** Statistical significance determined by a two-tailed Mann-Whitney test (ULF-STDP(+) n=1 mouse, n=3 cells; naiive n=5 mice, n=1-3 cells/mouse, p<0.05)

## Discussion

Here, we describe a novel approach for using STDP (spike-timing dependent plasticity) to reverse maladaptive hyperactivity of DMS D1-MSNs associated with AUD alcohol use disorder. This approach relies upon a method of delivering paired electromagnetic pulses through multiple electrodes at an ultra-low frequency (≤1Hz) to induce STDP onto D1-MSNs of the direct pathway. The results described here are consistent with an emerging body of literature suggesting that reducing alcohol-related D1-MSN hyperactivity or increasing D2-MSN hypoactivity reduces alcohol seeking and consumption ^17,20–29^. We tested the hypothesis that GPi/ACC ULF-STDP(+) will selectively decrease alcohol consumption and D1-MSN synaptic strength in a mouse model of AUD **(Figure 7)**. By applying ULF-STDP stimulation to the ACC and GPi, we expect to enable selective bidirectional gain control of glutamatergic ACC efferents to D1-MSNs (direct pathway) **(Figure 1D)**. Repeated phase-locked stimulation of the presynaptic axons in the cingulum bundle projecting from the ACC and the postsynaptic D1-MSN axons projecting to the GPi will avoid plasticity induction in D2-MSNs, which project to the GPe and will limit plasticity induction in other spatially distant brain regions.

**Figure 7:**
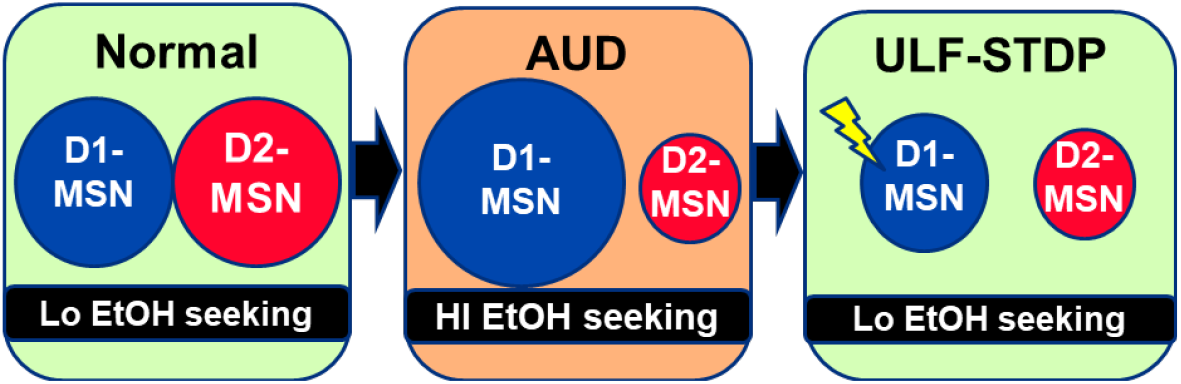
ULF-STDP(+) reduces DMS D1-MSN hyperactivity and alcohol consumption. Circle size represents the strength of corticostriatal glutamatergic synapses.

Our data show that ULF-STDP(+) decreases alcohol consumption without influencing food or water consumption. While the alcohol-specific effects of ULF-STDP(+) are promising, care must be taken not to induce hypoactivity of the direct pathway, as undesirable reductions in general locomotor activity^62^ and goal directed behavior^63^ have been reported. Further studies must be conducted to optimize ULF-STDP(+) dosing and re-dosing parameters (i.e. stimulation interval, stimulation conditions, etc.) in such a manner that maximizes reductions in alcohol consumption without negatively affecting general motor or reward-related behavior.

Assessment of neurological function at multiple scales is paramount to establish a causal functional role of a synaptic neuroplasticity induction protocol such as ULF-STDP in altering neural network function, and ultimately influencing animal behavior. We demonstrated that reduction in alcohol consumption mediated by ULF-STDP(+) is accompanied by neurological adaptations at multiple scales, including reductions in mean firing rate within the DMS **(Figure 4)**, decreases in evoked peak potentials **(Figure 5)**, and decreases in amplitude of D1-MSN mEPSCs **(Figure 6)**. Multiple factors influence the direction and magnitude of neuroplasticity induced by STDP protocols, including the relative timing of presynaptic and postsynaptic depolarizations, the number of pulse pairings, the cell types involved in the synapse, behavioral contexts, and extracellular levels of neuromodulators such as dopamine

Initial investigations into the STDP biophysical phenomenon were performed in cultured hippocampal cells where it was documented that repetitive postsynaptic spiking approximately 20 msec after presynaptic activation resulted in synaptic LTP while reversing the relative pre-post timing leads to LTD ^55^. Additional studies demonstrate that LTP and LTD occur in the corticostriatal synapse with similar pre-post activation timings ^67^. Conversely, it has also been repeatedly demonstrated that in the corticostriatal synapse, presynaptic depolarization prior to postsynaptic depolarization causes LTD ^68,69^. While our STDP findings may appear to disagree with prior studies, there are important methodological differences which explain these seemingly contradictory results. Namely, in Figure 3, ULF-STDP is delivered while the animal is awake and during a state of acute withdrawal. Acute withdrawal is associated with a decrease in extracellular dopamine ^70^, which biases STDP to produce LTD in D1-MSNs and LTP in D2-MSNs regardless of whether a pre- or post-synaptic action potential occurs first^57^. Additionally, it is important to note that the direct and indirect pathways are largely but not entirely comprised of D1-MSNs and D2-MSNs^71^, and the exact contribution of these D2-MSNs which project to the GPi is still unknown. To avoid the theoretical risk of de-potentiating the sparse GPi-projecting D2-MSNs, we delivered the postsynaptic stimulation prior to the presynaptic stimulation. Our ongoing studies are characterizing synaptic effects of ULF-STDP on D2-MSNs. Furthermore, we will explore whether alcohol consumption can also be reduced with a stimulation protocol in which the presynaptic cortical neurons are stimulated prior to the postsynaptic neurons. Future work must be performed to confirm the ULF-STDP(+) mechanism of action *in vivo*, which is expected to be dependent on mGluR5, dopamine, and CB-1 signaling^56,57,72^.

There are distinct advantages to move away from traditional high frequency DBS and towards pulse sequences designed to treat maladaptive neuroplasticity. First among these advantages is enhanced control of specific circuits of types of cells. While others have previously employed DBS to enable cell-specific reversal of drug-related maladaptive striatal plasticity^73,74^, our approach enables control of synaptic strength of D1-MSNs without pharmacological modulation or genetic modification through the use of only STDP-based DBS guided by neural pathway-based electrode placement. Reliance on electrical stimulation alone is paramount, as it obviates problems associated with clinical adoption of genetic modulation required for optogenetics and chemogenetics while avoiding undesirable off-target effects of pharmacological tools like D1-dopamine receptor antagonists^75^. Furthermore, this study represents a new direction for a clinical therapy like DBS in which the pulse sequence is designed to purposefully correct dysfunction associated with maladaptive neuroplasticity rather than treating symptoms in an acute manner. Additionally, low frequency and low duty cycle DBS confers the advantage of lower power requirements; resulting in improvement of battery life if delivered through implantable pulse generators and avoiding the need for surgical battery replacement which comes with additional costs and risk of infection.

Taken together, our results suggest that the development of novel neuromodulation approaches based on reversing maladaptive neuroplasticity underlying symptom-specific circuitopathies may improve existing invasive neuromodulation therapies and allow for expansion to new indications. ULF-STDP allows for network-level gain control in deep brain structures previously unachievable with existing clinical neuromodulation approaches. In turn, this may enable a new class of neuromodulation therapies for treatment of a wide range of disorders associated with maladaptive plasticity of the cortico-striato-pallidal pathway beyond AUD, such as Tourette’s syndrome ^76^, Obsessive Compulsive Disorder (OCD) ^77,78^, Schizophrenia ^79^, Parkinson’s Disease (PD) ^80–82^, and Manic Depression ^83^.

## Supplementary figures

**S1:**
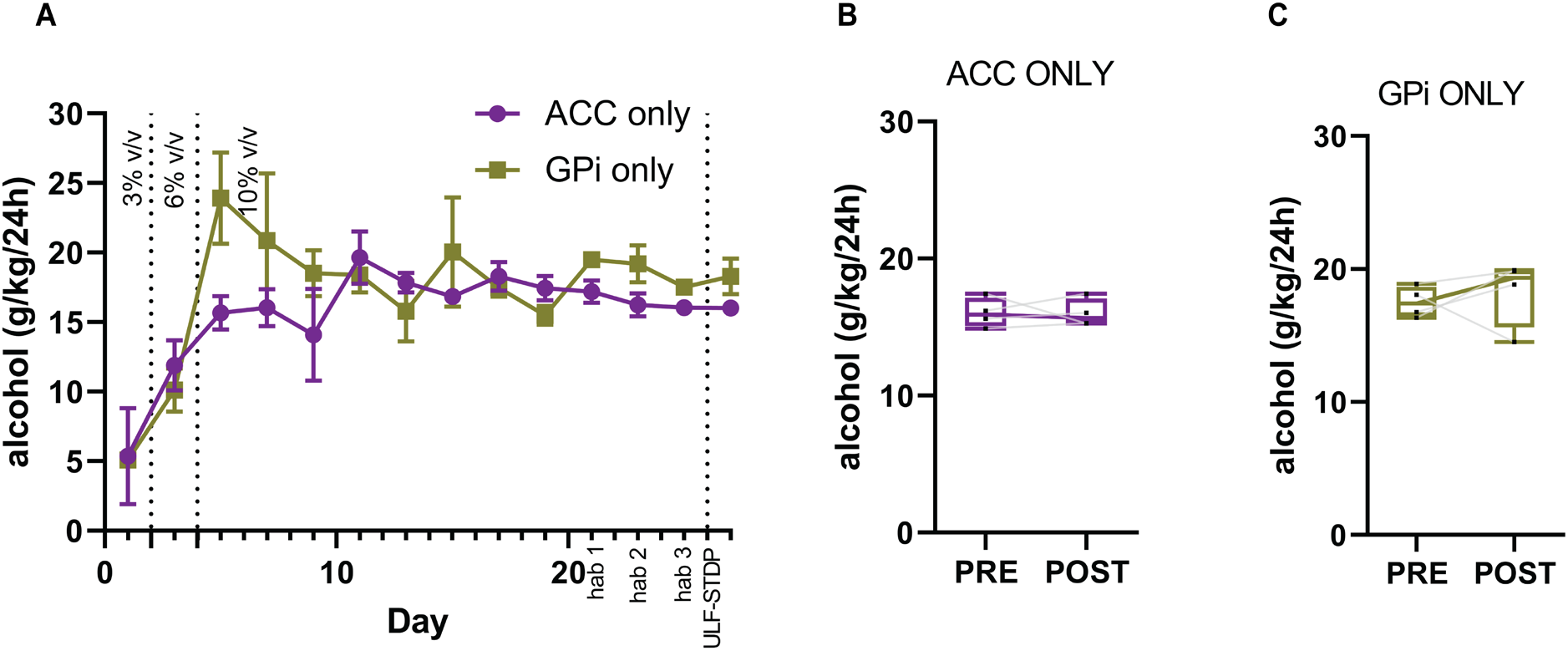
ULF-STDP protocol applied to either the ACC or GPi electrode alone does not alter alcohol consumption in a two-bottle choice model. In mice displaying high alcohol preference, ULF-STDP was applied to a single electrode in either the ACC (magenta) or GPi (gold) **(A)**. Alcohol consumption in g/kg/24h period before ULF-STDP (PRE) and 24 hours after ULF-STDP (POST) for groups in which ULF-STDP(+) was delivered to only the ACC **(B)** and ULF-STDP(+) GPi **(C).** Statistical significance of comparison of alcohol consumption immediately before and after ULF-STDP was determined using a paired Student’s t-test, p<0.05 (n=15). Error bars represent standard error. Box and whisker plots in B-C represent median, 95% CI, and max/minimum where black dots and gray lines represent individual animals.

**S2:**
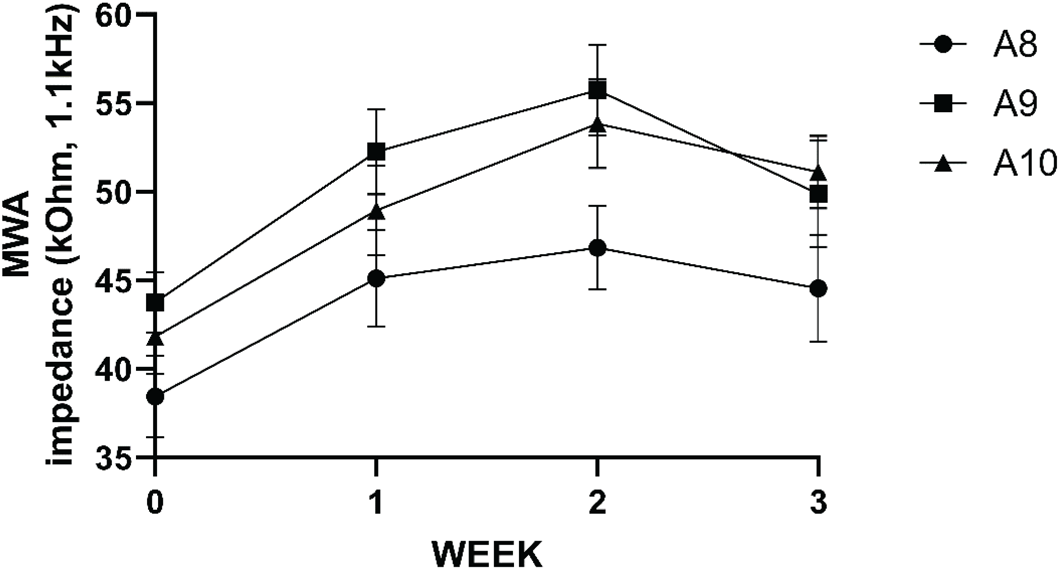
**Microwire array impedances** measured via TDT PZ5 neurodigitizer amplifier. Data are displayed as mean with error bars representing standard error of impedance values of all electrodes for individual animals identified as A8, A9, and A10.

**S3:**
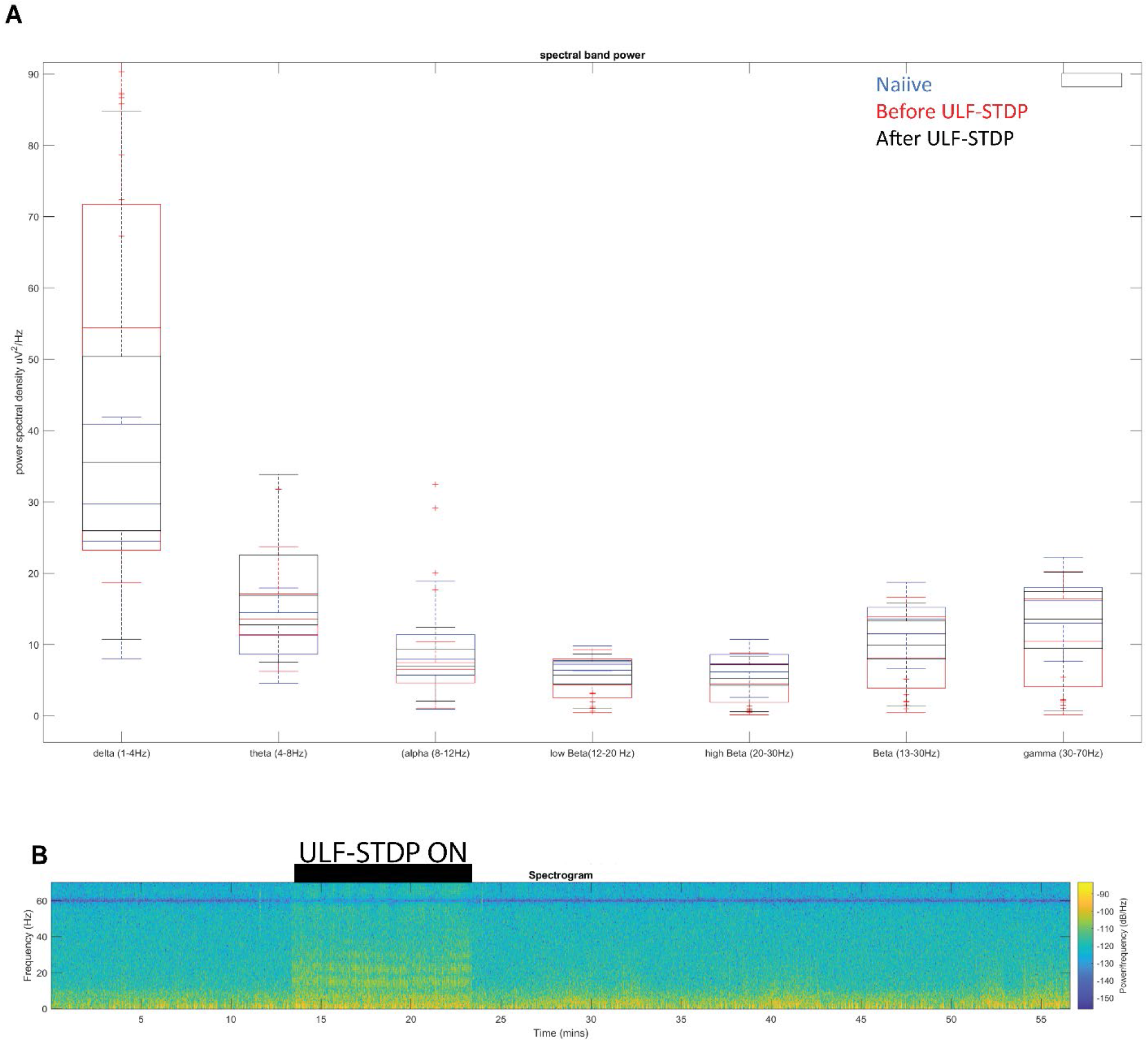
**Local field potential** band power analysis showing representative power spectral density(A) and spectrogram (B) before and after ULF-STDP (+)

